# Harmonization of resting-state functional MRI data across multiple imaging sites via the separation of site differences into sampling bias and measurement bias

**DOI:** 10.1101/440875

**Authors:** Ayumu Yamashita, Noriaki Yahata, Takashi Itahashi, Giuseppe Lisi, Takashi Yamada, Naho Ichikawa, Masahiro Takamura, Yujiro Yoshihara, Akira Kunimatsu, Naohiro Okada, Hirotaka Yamagata, Koji Matsuo, Ryuichiro Hashimoto, Go Okada, Yuki Sakai, Jun Morimoto, Jin Narumoto, Yasuhiro Shimada, Kiyoto Kasai, Nobumasa Kato, Hidehiko Takahashi, Yasumasa Okamoto, Saori C Tanaka, Mitsuo Kawato, Okito Yamashita, Hiroshi Imamizu

**Author notes:** Correspondence (H.I.), (O.Y.), or (A.Y.).

## Abstract

When collecting large neuroimaging data associated with psychiatric disorders, images must be acquired from multiple sites because of the limited capacity of a single site. However, site differences represent the greatest barrier when acquiring multi-site neuroimaging data. We utilized a traveling-subject dataset in conjunction with a multi-site, multi-disorder dataset to demonstrate that site differences are composed of biological sampling bias and engineering measurement bias. Effects on resting-state functional MRI connectivity because of both bias types were greater than or equal to those because of psychiatric disorders. Furthermore, our findings indicated that each site can sample only from among a subpopulation of participants. This result suggests that it is essential to collect large neuroimaging data from as many sites as possible to appropriately estimate the distribution of the grand population. Finally, we developed a novel harmonization method that removed only the measurement bias by using traveling-subject dataset and achieved the reduction of the measurement bias by 29% and the improvement of the signal to noise ratios by 40%.

## Introduction

Acquiring and sharing large neuroimaging data have recently become critical for bridging the gap between basic neuroscience research and clinical applications such as the diagnosis and treatment of psychiatric disorders (Human Connectome Project (HCP) [1], [https://www.humanconnectomeproject.org/]; Human Brain Project [http://www.humanbrainproject.eu/en/]; UK Biobank [http://www.ukbiobank.ac.uk/]; and Strategic Research Program for Brain Sciences (SRPBS) [2] [http://www.amed.go.jp/program/list/01/04/001_nopro.html]) [3–5]. When collecting large data associated with psychiatric disorders, it is necessary to acquire images from multiple sites because it is nearly impossible for a single site to collect large neuroimaging data (Connectomes Related to Human Disease (CRHD), [http://www.humanconnectome.org/disease-studies]; Autism Brain Imaging Data Exchange (ABIDE); and SRPBS) [2, 6–8]. In 2013, the Japan Agency for Medical Research and Development (AMED) organized the Decoded Neurofeedback (DecNef) Project. The project determined the unified imaging protocol on 28th February 2014 (http://www.cns.atr.jp/rs-fmri-protocol-2) and have collected multisite resting-state functional magnetic resonance imaging (rs-fMRI) data using twelve scanners across eight research institutes for recent five years. The collected dataset encompasses 2,239 samples and five disorders and is publicly shared through the SRPBS multisite multi-disorder database (https://bicr-resource.atr.jp/decnefpro/). This project has enabled the identification of resting-state functional connectivity (rs-fcMRI)-based biomarkers of several psychiatric disorders that can be generalized to completely independent cohorts [2, 8–10]. However, multisite dataset with multiple disorders raises difficult problems never included in a single-site based dataset of healthy population (e.g., HCP and UK Biobank). That is, our experience in the SRPBS database demonstrated difficulty in fully control of scanner type, imaging protocol, patient demographics [10–13] even if the unified protocol is determined. Moreover, there often exists unpredictable difference in participant population among sites. Therefore, researchers must work with heterogeneous neuroimaging data. In particular, site differences represent the greatest barrier when extracting disease factors by applying machine-learning techniques to such heterogeneous data [14] because disease factors tend to be confounded with site factors [2, 8, 10–13, 15]. This confounding occurs because a single site (or hospital) is apt to sample only a few types of psychiatric disorders (e.g., primarily schizophrenia from site A and primarily autism spectrum disorder from site B). To properly manage such heterogeneous data, it is necessary to harmonize the data among the sites [16–19]. Moreover, a deeper understanding of these site differences is essential for efficient harmonization of the data.

Site differences essentially consist of two types of biases: engineering bias (i.e., measurement bias) and biological bias (i.e., sampling bias). Measurement bias includes differences in the properties of MRI scanners such as imaging parameters, field strength, MRI manufacturers, and scanner models, whereas sampling bias refers to differences in participant groups among sites. Previous studies have investigated the effect of measurement bias on resting-state functional connectivity by using a traveling-subject design [20] wherein multiple participants travel to multiple sites for the assessment of measurement bias [7]. By contrast, researchers to date have only speculated with regard to sampling bias. For example, differences in the clinical characteristics of patients examined at different sites are presumed to underlie the stagnant accuracy of certain biomarkers, even after combining the data from multiple sites [12]. Furthermore, to our knowledge, no study has mathematically defined sampling bias or conducted quantitative analyses of its effect size, which is likely because the decomposition of site differences into measurement bias and sampling bias is a complex process. To achieve this aim, we combined a separate traveling-subject rs-fMRI dataset with the SRPBS multi-disorder dataset. Simultaneous analysis of the datasets enabled us to divide site differences into measurement bias and sampling bias and to quantitatively compare their effect sizes on resting-state functional connectivity with those of psychiatric disorders.

Furthermore, our detailed analysis of measurement and sampling biases enabled us to investigate the origin of each bias in multisite datasets for the first time. For measurement bias, we quantitatively compared the magnitude of the effects among the different imaging parameters, fMRI manufacturers, and number of coils in each fMRI scanner. We further examined two alternative hypotheses regarding the mechanisms underlying sampling bias: one hypothesis assumes that each site samples subjects from a common population. In this situation, sampling bias occurs because of the random sampling of subjects, which results in incidental differences in the patients’ characteristics among the sites. The second hypothesis assumes that each site samples subjects from different subpopulations. In this situation, sampling bias occurs because of sampling from subpopulations with different characteristics. For example, assume multiple sites plan to collect data from the same population of patients with major depressive disorder. Subtypes of major depressive disorder exist within the population such as atypical depression and melancholic depression [21, 22]; therefore, one subpopulation may contain a large proportion of patients with atypical depression, whereas another subpopulation may contain a large proportion of patients with melancholic depression. Therefore, in some instances, atypical depression may be more frequent among patients at site A, whereas melancholic depression may be more frequent among patients at site B. The basic protocol for collecting large-scale datasets differ between these two hypotheses; thus, it is necessary to determine the hypothesis that most appropriately reflects the characteristics of the SRPBS dataset. In the former situation, one would simply need to collect data from a large number of subjects, even with a small number of sites. In the latter situation, a larger number of sites would be required to obtain truly representative data.

To overcome these limitations associated with site differences, we developed a novel harmonization method that enabled us to subtract only the measurement bias by using a traveling-subject dataset. We investigated that how much our proposed method could reduce the measurement bias and could improve the signal to noise ratio. We compared its performance to those of other commonly used harmonization methods. All data utilized in this study can be downloaded publicly from the DecNef Project Brain Data Repository at https://bicr-resource.atr.jp/decnefpro/.

## Results

### Datasets

We used two rs-fMRI datasets: the (1) SRPBS multi-disorder dataset, (2) a traveling-subject dataset.

#### SRPBS multi-disorder dataset

This dataset included patients with five different disorders and healthy controls (HCs) who were examined at nine sites belonging to eight research institutions. A total of 805 participants were included: 482 HCs from nine sites, 161 patients with major depressive disorder (MDD) from five sites, 49 patients with autism spectrum disorder (ASD) from one site, 65 patients with obsessive-compulsive disorder (OCD) from one site, and 48 patients with schizophrenia (SCZ) from three sites (Supplementary Table 1). The rs-fMRI data were acquired using a unified imaging protocol at all but three sites (Supplementary Table 2; http://www.cns.atr.jp/rs-fmri-protocol-2/). Site differences in this dataset included both measurement and sampling biases (Fig. 1a). For bias estimation, we only used data obtained using the unified protocol. (Patients with OCD were not scanned using this unified protocol; therefore, the disorder factor could not be estimated for OCD.)

**Figure 1:**
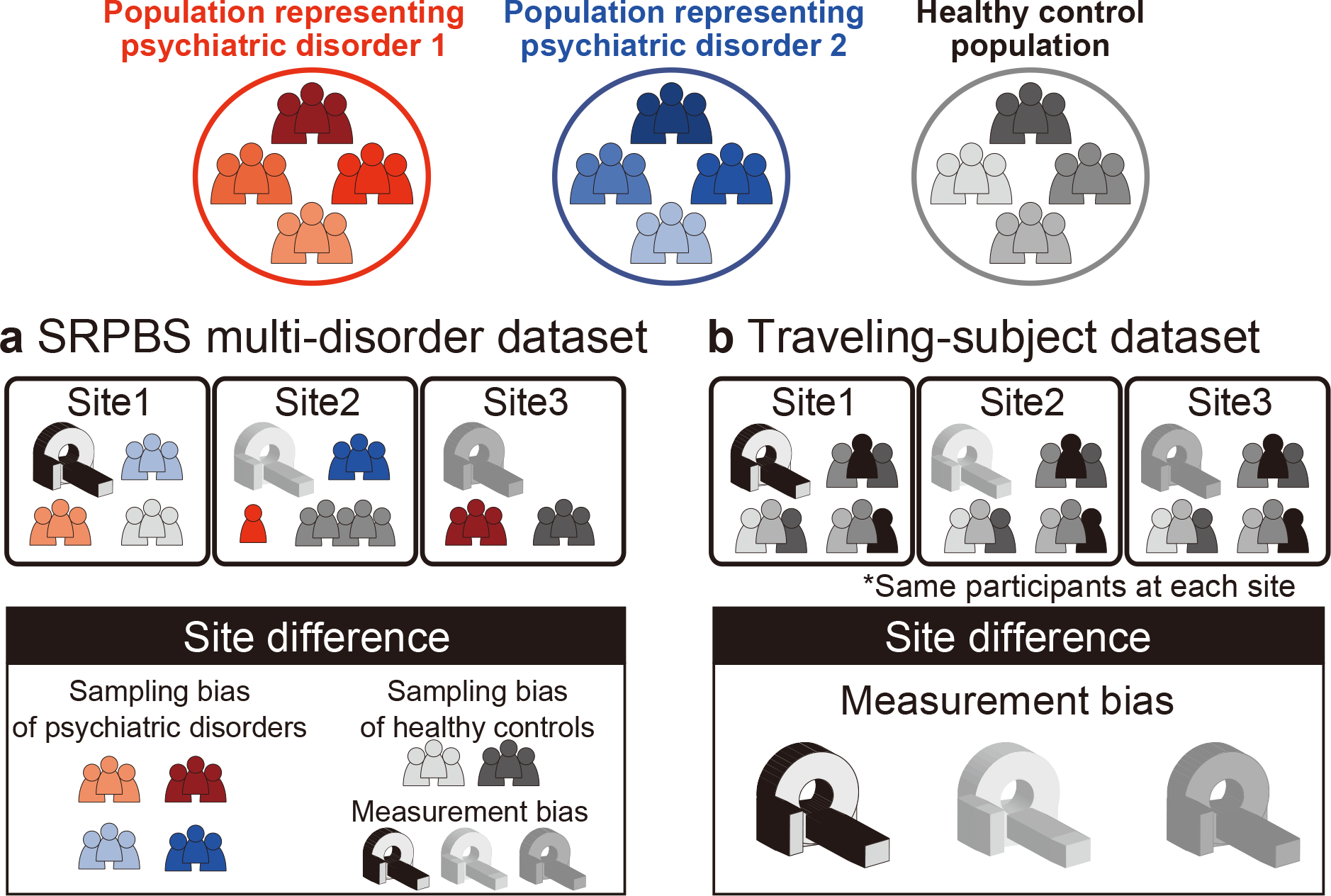
Schematic examples illustrating the two main datasets. (a) The SRPBS multi-disorder dataset includes patients with psychiatric disorders and healthy controls. The number of patients and scanner types differed among sites. Thus, site differences consist of sampling bias and measurement bias. (b) The traveling-subject dataset includes only healthy controls, and the participants were the same across all sites. Thus, site differences consist of measurement bias only. SRPBS: Strategic Research Program for Brain Sciences.

#### Traveling-subject dataset

We acquired a traveling-subject dataset to estimate measurement bias across sites in the SRPBS dataset. Nine healthy participants (all men; age range: 24–32 years; mean age: 27±2.6 years) were scanned at each of 12 sites, which included the nine sites in the SRPBS dataset, and produced a total of 411 scan sessions (see “Participants” in the Methods section). Although we had attempted to acquire this dataset using the same imaging protocol as that in the SRPBS multi-disorder dataset, there were some differences in the imaging protocol across sites because of limitations in parameter settings or the scanning conventions of each site (Supplementary Table 3). There were two phase-encoding directions (P→A and A→P), three MRI manufacturers (Siemens, GE, and Philips), four numbers of coil channels (8, 12, 24, and 32), and seven scanner types (TimTrio, Verio, Skyra, Spectra, MR750W, SignaHDxt, and Achieva). Site differences in this dataset included measurement bias only as the same nine participants were scanned across the 12 sites (Fig. 1b).

### Visualization of site differences and disorder effects

We first visualized the site differences and disorder effects in the SRPBS multi-disorder rs-fcMRI dataset while maintaining its quantitative properties by using a principal component analysis (PCA)—an unsupervised dimension reduction method. Functional connectivity was calculated as the temporal correlation of rs-fMRI blood-oxygen-level dependent (BOLD) signals between two brain regions for each participant. There are some candidates for the measure of functional connectivity such as the tangent method and partial correlation [11, 23]; however, we used Pearson’s correlation coefficients because they have been the most commonly used values in previous studies. Functional connectivity was defined based on a functional brain atlas consisting of 268 nodes (i.e., regions) covering the whole brain, which has been widely utilized in previous studies [20, 24–26]. The Fisher’s *z*-transformed Pearson’s correlation coefficients between the preprocessed BOLD signal time courses of each possible pair of nodes were calculated and used to construct 268 × 268 symmetrical connectivity matrices in which each element represents a connection strength, or edge, between two nodes. We used 35,778 connectivity values [i.e., (268 × 267)/2] of the lower triangular matrix of the connectivity matrix. All participant data in the SRPBS multi-disorder dataset were plotted on two axes consisting of the first two principal components (Fig. 2, small, light-colored symbols). The averages of the HCs within individual sites and the averages of individual psychiatric or developmental disorders are presented as dark-colored symbols in Fig. 2. There was a clear separation of the Hiroshima University Hospital (HUH) site for principal component 1, which explained most of the variance in the data. Furthermore, there were no differences between the differences of the sites and the disorder factors. Patients with ASD were only scanned at the Showa University (SWA) site; therefore, the averages for patients with ASD (▲) and HCs (blue ●) scanned at this site were projected to nearly identical positions (Fig. 2).

**Figure 2:**
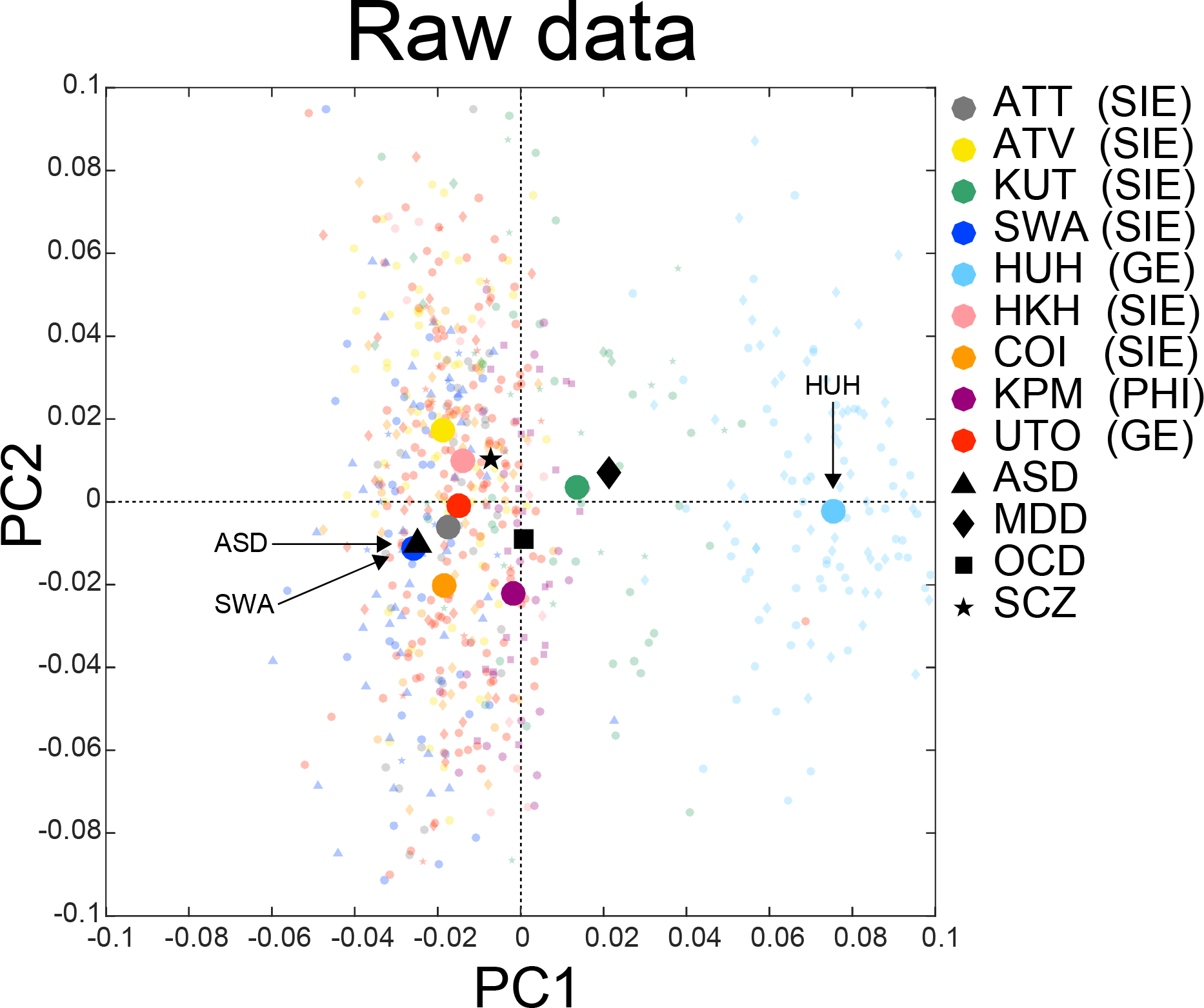
PCA dimension reduction in the SRPBS multi-disorder dataset. All participants in the SRPBS multi-disorder dataset projected into the first two principal components (PCs), as indicated by small, light-colored markers. The average across all healthy controls in each site and the average within each psychiatric disorder are depicted as dark-colored makers. The color of the marker represents the site, while the shape represents the psychiatric disorder. PCA: principal component analysis; SRPBS: Strategic Research Program for Brain Sciences; ATT: Siemens TimTrio scanner at Advanced Telecommunications Research Institute International; ATV: Siemens Verio scanner at Advanced Telecommunications Research Institute International; KUT: Siemens TimTrio scanner at Kyoto University; SWA: Showa University; HUH: Hiroshima University Hospital; HKH: Hiroshima Kajikawa Hospital; COI: Center of Innovation in Hiroshima University; KPM: Kyoto Prefectural University of Medicine; UTO: University of Tokyo; ASD: Autism Spectrum Disorder. MDD: Major Depressive Disorder. OCD: Obsessive Compulsive Disorder. SCZ: Schizophrenia. SIE: Siemens fMRI. GE: GE fMRI. PHI: Philips fMRI.

### Bias estimation

To quantitatively investigate the site differences in the rs-fcMRI data, we identified measurement biases, sampling biases, and disorder factors. We defined measurement bias for each site as a deviation of the correlation value for each functional connection from its average across all sites. We assumed that the sampling biases of the HCs and patients with psychiatric disorders differed from one another. Therefore, we calculated the sampling biases for each site separately for HCs and patients with each disorder. Disorder factors were defined as deviations from the HC values. Sampling biases were estimated for patients with MDD and SCZ because only these patients were sampled at multiple sites. Disorder factors were estimated for MDD, SCZ, and ASD because patients with OCD were not scanned using the unified protocol.

It is difficult to separate site differences into measurement bias and sampling bias using only the SRPBS multi-disorder dataset because the two types of bias covaried across sites. Different samples (i.e., participants) were scanned using different parameters (i.e., scanners and imaging protocols). In contrast, the traveling-subject dataset included only measurement bias because the participants were fixed. By combining the traveling-subject dataset with the SRPBS multi-disorder dataset, we simultaneously estimated measurement bias and sampling bias as different factors affected by different sites. We utilized a linear mixed-effects model to assess the effects of both types of bias and disorder factors on functional connectivity, as follows.

#### Linear mixed-effects model for the SRPBS multi-disorder dataset

In this model, the connectivity values of each participant in the SRPBS multi-disorder dataset were composed of fixed and random effects. Fixed effects included the sum of the average correlation values across all participants and all sites at baseline, the measurement bias, the sampling bias, and the disorder factors. The combined effect of participant factors (i.e., individual difference) and scan-to-scan variations was regarded as the random effect (see “Estimation of biases and factors” in the Methods section).

#### Linear mixed-effects model for the traveling-subject dataset

In this model, the connectivity values of each participant for a specific scan in the traveling-subject dataset were composed of fixed and random effects. Fixed effects included the sum of the average correlation values across all participants and all sites, participant factors, and measurement bias. Scan-to-scan variation was regarded as the random effect. For each participant, we defined the participant factor as the deviation of connectivity values from the average across all participants.

We estimated all biases and factors by simultaneously fitting the aforementioned two regression models to the functional connectivity values of the two different datasets. For this regression analysis, we used data from participants scanned using a unified imaging protocol in the SRPBS multi-disorder dataset and from all participants in the traveling-subject dataset. In summary, each bias or each factor was estimated as a vector that included a dimension reflecting the number of connectivity values (i.e., 35,778). Vectors included in our further analyses are those for measurement bias at 12 sites, sampling bias of HCs at six sites, sampling bias for patients with MDD at three sites, sampling bias for patients with SCZ at three sites, participant factors of nine traveling-subjects, and disorder factors for MDD, SCZ, and ASD.

### Quantification of site differences

To quantitatively evaluate the effect of measurement and sampling biases on functional connectivity, we compared the magnitudes of both types of bias with the magnitudes of psychiatric disorders and participant factors. For this purpose, we investigated the magnitude distribution of both biases, as well as the effects of psychiatric disorders and participant factors on functional connectivity overall 35,778 elements in a 35,778-dimensional vector to see how many functional connectivities were largely affected (Fig. 3a: the *x*-axis shows the magnitude as Fisher’s *z*-transformed Pearson’s correlation coefficients, while the *y*-axis shows the density of the number of connectivities). Figure 3b shows the same data, except the *y*-axis represents the log-transformed number of connectivities for better visualization of small values. These distributions show that, on average, connectivity was unaffected by either type of bias or by each factor because the averages of each distribution were nearly 0. However, there were significant differences among biases and factors for larger magnitudes near the tails of their distributions. For example, the number of connectivities, which was largely affected (i.e., a magnitude larger than 0.2), was more than 100 for the participant factor, approximately 100 for measurement bias, and nearly 0 for all sampling biases, as well as all disorder factors.

**Figure 3:**
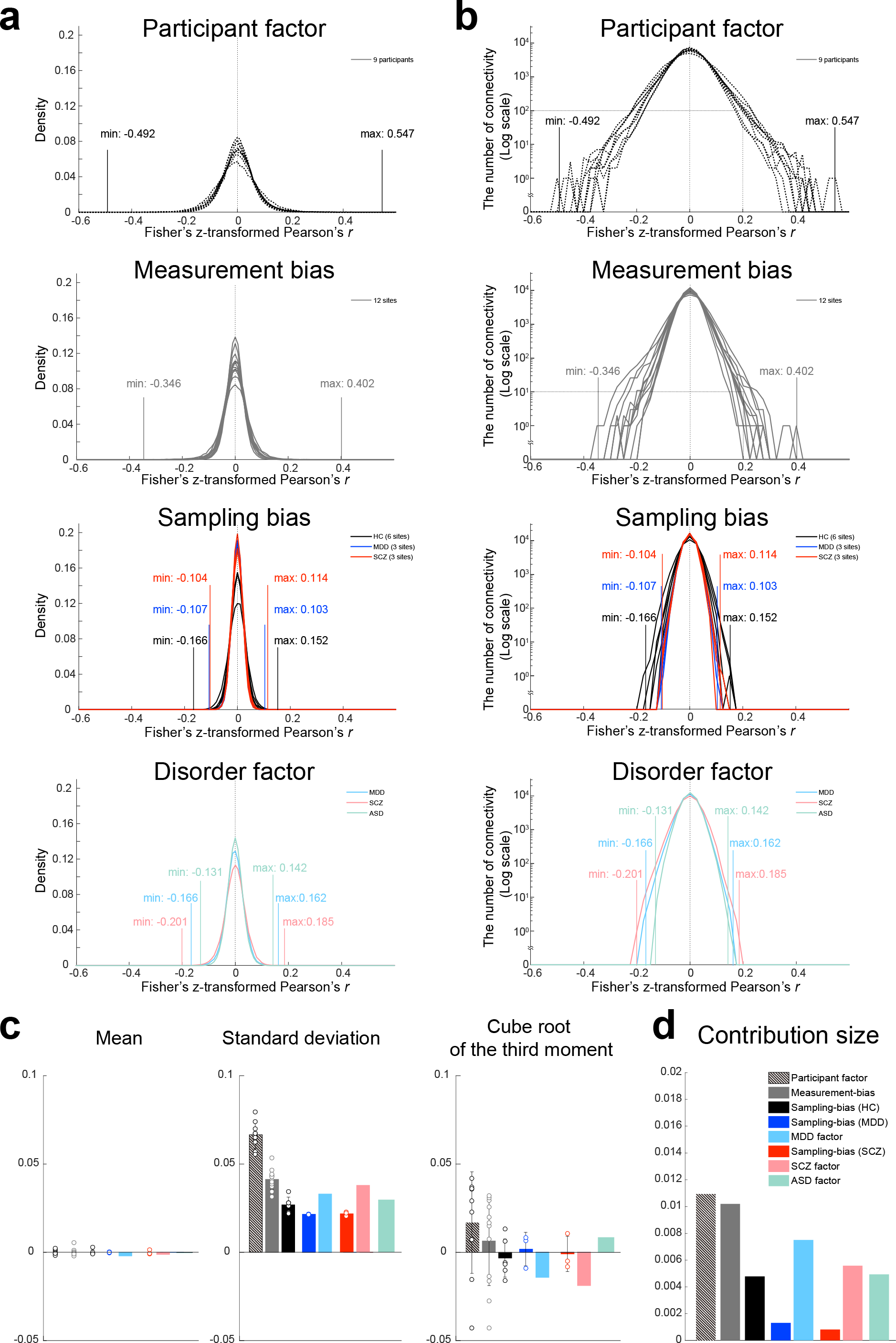
Distributions and statistics for each type of bias and each factor. (a, b) The distribution of the effects of each bias and each factor on functional connectivity vectors. Functional connectivity was measured based on Fisher’s z-transformed Pearson’s correlation coefficients. The *x*-axis represents the effect size of the Fisher’s z-transformed Pearson’s correlation coefficients. In (a) and (b), the *y*-axis represents the density of connectivity and the log-transformed the number of connections, respectively. Each line represents one participant or one site. (c) The means, standard deviations, and third moments standardized to the same scale on the vertical axis (i.e., cube root) for each type of bias and each factor. Bars represent the average value, while the error bars represent the standard deviation across sites or participants. Each data point represents one participant or one site. (d) Contribution size of each bias and each factor. HC: healthy controls; SCZ: schizophrenia; MDD: major depressive disorder; ASD: autism spectrum disorder.

To quantitatively summarize the effect of each factor, we calculated the first, second, and third statistical moments of each histogram (Fig. 3c). Based on the mean values and the cube roots of the third moments, all distributions could be approximated as bilaterally symmetric with a mean of zero. Thus, distributions with larger squared roots of the second moments (standard deviations) affect more connectivities with larger effect sizes. The value of the standard deviation was largest for the participant factor (0.0662), followed by these values for the measurement bias (0.0411), the SCZ factor (0.0377), the MDD factor (0.0328), the ASD factor (0.0297), the sampling bias for HCs (0.0267), sampling bias for patients with SCZ (0.0217), and sampling bias for patients with MDD (0.0214). To compare the sizes of the standard deviation between participant factors and measurement bias, we evaluated the variance of each distribution. All pairs of variances were analyzed using Ansari–Bradley tests. Our findings indicated that all variances of the participant factors were significantly larger than all variances of the measurement biases (nine participant factors × 12 measurement biases = 108 pairs; *W*\*: mean = −59.80, max = −116.81, min = −3.69; *p* value after Bonferroni correction: max = 0.011, min = 0, *n* = 35,778). In addition, the variances of 10 of 12 measurement biases were significantly larger than the variance of the MDD factor, the variances of seven of 12 measurement biases were significantly larger than the variance of the SCZ factor, and the variances of all measurement biases were significantly larger than the variance of the MDD factor (Supplementary Table 8). Furthermore, we plotted fractions of the data variance determined using the aforementioned factors (i.e., contribution size) in our linear mixed-effects model (Fig. 3d; see “Analysis of contribution size” in the Methods section). The results were consistent with the analysis of the standard deviation (Fig. 3c, middle). These results indicate that the effect size of measurement bias on functional connectivity is smaller than that of the participant factor but is mostly larger than those of the disorder factors, which suggest that measurement bias represents a serious limitation in research regarding psychiatric disorders. The largest variance in sampling bias was significantly larger than the variance of the MDD factor (Supplementary Table 9), whereas the smallest variance in sampling bias was one-half the size of the variance for disorder factors. These findings indicate that sampling bias also represents a major limitation in psychiatric research.

The standard deviation of the participant factor was approximately twice that for SCZ, MDD, and ASD; therefore, individual variability within the healthy population was much greater than that among patients with SCZ, MDD, or ASD when all functional connections were considered. Furthermore, the standard deviations of the measurement biases were mostly larger than those of the disorder factors, while the standard deviations of the sampling biases were comparable with those of the disorder factors. Such relationships make the development of rs-fcMRI-based classifiers of psychiatric or developmental disorders very challenging. Only when a small number of disorder-specific and site-independent abnormal functional connections can be selected from among a vast number does it become feasible to develop robust and generalizable classifiers across multiple sites [2, 8–10, 15].

### Brain regions contributing most to biases and associated factors

To evaluate the spatial distribution of the two types of bias and all factors in the whole brain, we utilized a previously described visualization method [27] to project connectivity information to anatomical regions of interest (ROIs). We first quantified the effect of a bias or a factor on each functional connectivity as the median of its absolute values across sites or across participants. Thus, we obtained 35,778 values, each of which was associated with one connectivity and represented the effect of a bias or factor on the connectivity. We then summarized these effects on connectivity for each ROI by averaging the values of all connectivities connected with the ROI (see “Spatial characteristics of measurement bias, sampling bias, and each factor in the brain” in the Methods section). The average value represents the extent the ROI contributes to the effect of a bias or factor. By repeating this procedure for each ROI and coding the averaged value based on the color of an ROI, we were able to visualize the relative contribution of individual ROIs to each bias or factor in the whole brain (Fig. 4). Consistent with the findings of previous studies, the effect of the participant factor was large for several ROIs in the cerebral cortex, especially in the prefrontal cortex, but small in the cerebellum and visual cortex [24]. The effect of measurement bias was large in inferior brain regions where functional images are differentially distorted depending on the phase-encoding direction [28, 29]. Connections involving the medial dorsal nucleus of the thalamus were also heavily affected by both MDD, SCZ and ASD. Effects of the MDD factor were observed in the dorsomedial prefrontal cortex and the superior temporal gyrus in which abnormalities have also been reported in previous studies [22, 30, 31]. Effects of the SCZ factor were observed in the left inferior parietal lobule, bilateral anterior cingulate cortices, and left middle frontal gyrus in which abnormalities have been reported in previous studies [32–34]. Effects of the ASD factor were observed in the putamen, the medial prefrontal cortex, and the right middle temporal gyrus in which abnormalities have also been reported in previous studies [10, 11, 35]. The effect of sampling bias for HCs was large in the inferior parietal lobule and the precuneus, both of which are involved in the default mode network and the middle frontal gyrus. Sampling bias for disorders was large in the medial dorsal nucleus of the thalamus, left dorsolateral prefrontal cortex, dorsomedial prefrontal cortex, and cerebellum for MDD [22]; and in the prefrontal cortex, cuneus, and cerebellum for SCZ [33].

**Figure 4:**
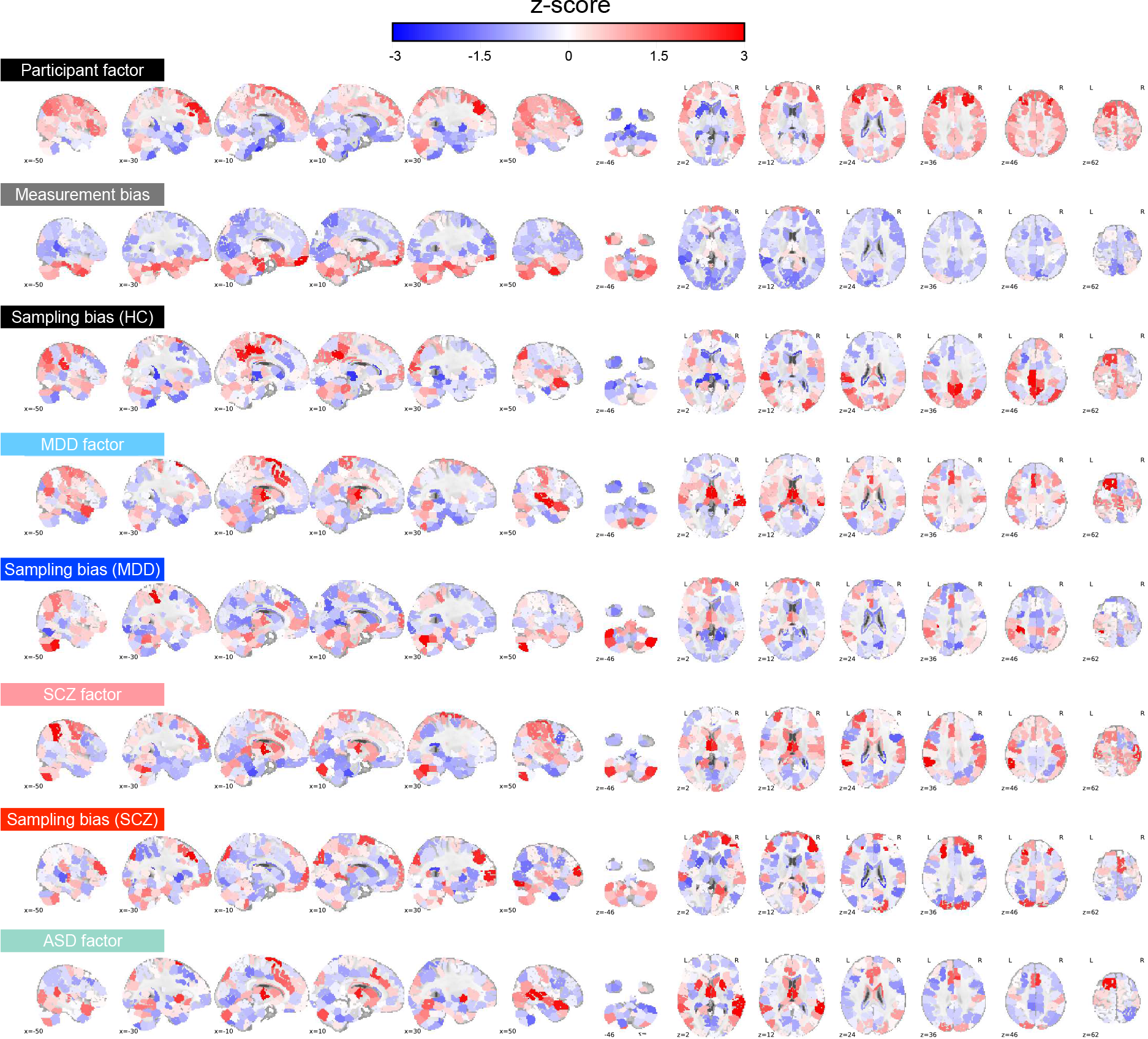
Spatial distribution of each type of bias and each factor in various brain regions. Mean effects of connectivity for all 268 ROIs. For each ROI, the mean effects of all functional connections associated with that ROI were calculated for each bias and each factor. Warmer (red) and cooler (blue) colors correspond to large and small effects, respectively. The magnitudes of the effects are normalized within each bias or each factor (*z*-score). ROI: region of interest; HC: healthy control; SCZ: schizophrenia; MDD: major depressive disorder; ASD: autism spectrum disorder.

### Characteristics of measurement bias

We next investigated the characteristics of measurement bias. We first examined whether similarities among the estimated measurement bias vectors for the 12 included sites reflect certain properties of MRI scanners such as phase-encoding direction, MRI manufacturer, coil type, and scanner type. We used hierarchical clustering analysis to discover clusters of similar patterns for measurement bias. This method has previously been used to distinguish subtypes of MDD, based on rs-fcMRI data [22]. As a result, the measurement biases of the 12 sites were divided into phase-encoding direction clusters at the first level (Fig. 5a). They were divided into fMRI manufacturer clusters at the second level, and further divided into coil type clusters, followed by scanner model clusters. Furthermore, we quantitatively verified the magnitude relationship among factors by using the same model to assess the contribution of each factor (Fig. 5b; see “Analysis of contribution size” in the Methods section). The contribution size was largest for the phase-encoding direction (0.0391), followed by the contribution sized for fMRI manufacturer (0.0318), coil type (0.0239), and scanner model (0.0152). These findings indicate that the main factor influencing measurement bias is the difference in the phase-encoding direction, followed by fMRI manufacturer, coil type, and scanner model, respectively.

**Figure 5:**
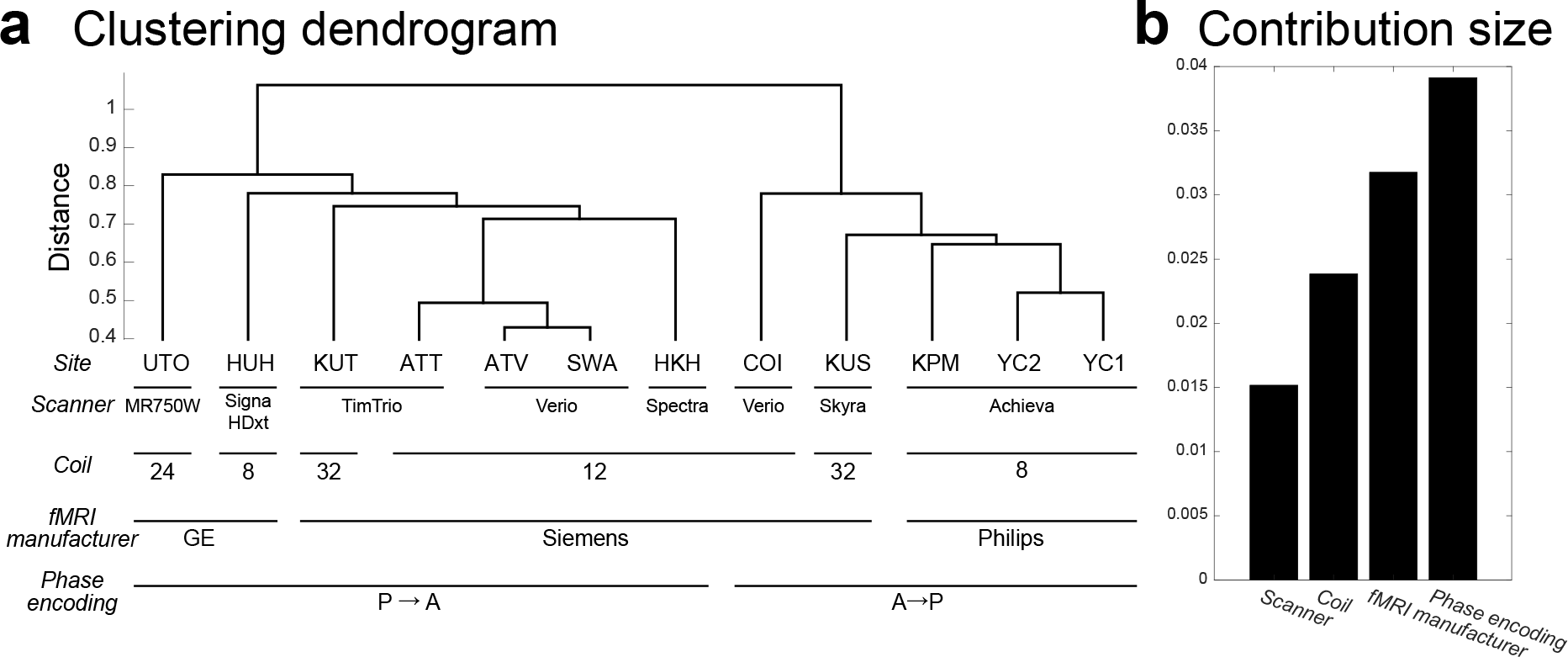
Clustering dendrogram for measurement bias. (a) The height of each linkage in the dendrogram represents the dissimilarity (1 - *r*) between the clusters joined by that link. (b) Contribution size of each factor. UTO: University of Tokyo; HUH: Hiroshima University Hospital; KUT: Siemens TimTrio scanner at Kyoto University; ATT: Siemens TimTrio scanner at Advanced Telecommunications Research Institute International; ATV: Siemens Verio scanner at Advanced Telecommunications Research Institute International; SWA: Showa University; HKH: Hiroshima Kajikawa Hospital; COI: Center of Innovation in Hiroshima University; KUS: Siemens Skyra scanner at Kyoto University; KPM: Kyoto Prefectural University of Medicine; YC1: Yaesu Clinic 1; YC2: Yaesu Clinic 2.

### Sampling bias is because of sampling from among a subpopulation

We investigated two alternative models for the mechanisms underlying sampling bias. In the “single-population model”, which assumes that participants are sampled from a common population (Fig. 6a), the functional connectivity values of each participant were generated from a Gaussian distribution (see “Comparison of models for sampling bias” in the Methods section). In the “different-subpopulation model,” which assumes that sampling bias occurs partly because participants are sampled from among a different subpopulation at each site (Fig. 6b), we assumed that the average of the subpopulation differed among sites and was generated from a Gaussian distribution. In addition, the functional connectivity values of each participant were generated from a Gaussian distribution, based on the average of the subpopulation at each site. It is necessary to determine which model is more suitable for collecting big data across multiple sites: If the former model is correct, then the data can be used to represent a population by increasing the number of participants, even if the number of sites is small. If the latter model is correct, data should be collected from many sites, as a single site does not represent the true grand population distribution, even with a very large sample size.

**Figure 6:**
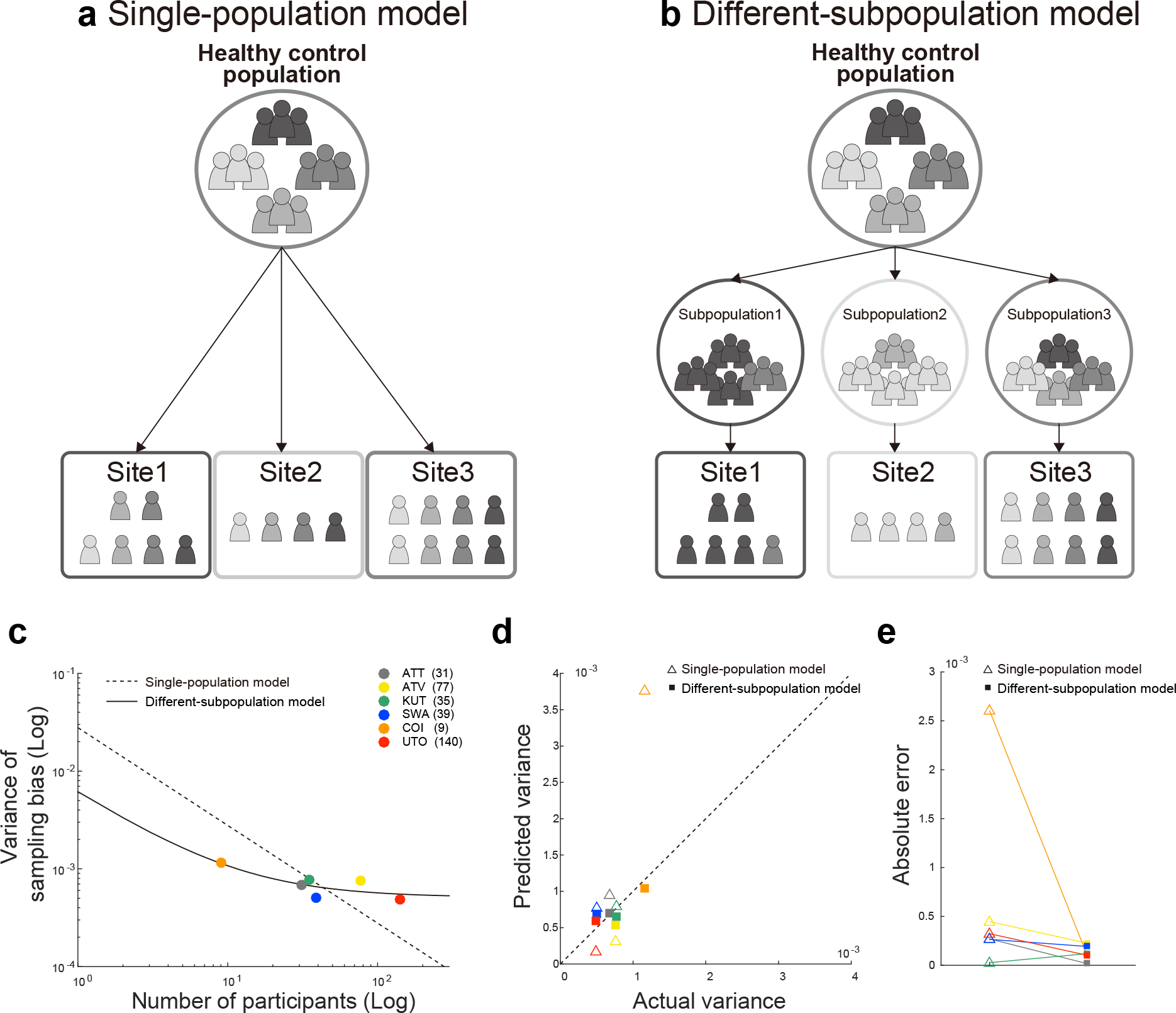
Comparison of the two models of sampling bias. Schematic examples illustrating the single-population (a) and different-subpopulation models and the results of model fitting (c). The *x*-axis represents the number of participants on a logarithmic scale, while the *y*-axis represents the variance of sampling bias on a logarithmic scale. The broken line represents the prediction of the single-population model, while the solid line represents the prediction of the different-subpopulation model. Each data point represents one site. (d) Results of the predictions determined by using each model. The *x*-axis represents the actual variance, while the *y*-axis represents the predicted variance. Open triangles correspond to the single-population model, while filled squares correspond to the different-subpopulation model. (e) Performance of prediction using the two models, based on the absolute error between the actual and predicted variance. UTO: University of Tokyo; COI: Center of Innovation in Hiroshima University; SWA: Showa University; KUT: Siemens TimTrio scanner at Kyoto University; ATT: Siemens TimTrio scanner at Advanced Telecommunications Research Institute International; ATV: Siemens Verio scanner at Advanced Telecommunications Research Institute International.

For each model, we first investigated how the number of participants at each site determined the effect of sampling bias on functional connectivity. We measured the magnitude of the effect, based on the variance values for sampling bias across functional connectivity (see the “Quantification of site differences” section). We used variance instead of the standard deviation to simplify the statistical analysis, although there is essentially no difference based on which value is used. We theorized that each model represents a different relationship between the number of participants and the variance of sampling bias. Therefore, we investigated which model best represents the actual relationships in our data by comparing the corrected Akaike information criterion (AICc) [36, 37] and Bayesian information criterion (BIC). Moreover, we performed leave-one-site-out cross-validation evaluations of predictive performance in which all but one site was used to construct the model and the variance of the sampling bias was predicted for the remaining site. We then compared the predictive performances between the two models. Our results indicated that the different-subpopulation model provided a better fit for our data than the single-population model (Fig. 6c; different subpopulation model: AICc = −108.80 and BIC = −113.22; single-population model: AICc = −96.71 and BIC = −97.92). Furthermore, the predictive performance was significantly higher for the different-subpopulation model than for the single-population model (one-tailed Wilcoxon signed-rank test applied to absolute errors: Z = 1.67, *p* = .0469, *n* = 6; Figs. 6d and 6e). This result indicates that sampling bias is not only caused by random sampling from a single grand population, depending on the number of participants among sites, but also by sampling from among different subpopulations. Sampling biases thus represent a major limitation in attempting to estimate a true single distribution of HC or patient data based on measurements obtained from a finite number of sites and participants.

### Visualization of the effect of harmonization

We next developed a novel harmonization method that enabled us to subtract only the measurement bias using the traveling-subject dataset. Using a linear mixed-effects model, we estimated the measurement bias separately from sampling bias (see the “Bias estimation” in the Methods section). Thus, we could remove the measurement bias from the SRPBS multi-disorder dataset (i.e., traveling-subject method, see “Traveling-subject harmonization” in the Methods section). To visualize the effects of the harmonization process, we plotted the data after subtracting only the measurement bias from the SRPBS multi-disorder dataset as described in the “Visualization of site differences and disorder effects” section (Fig. 7). Relative to the data reported in Fig. 2, which reflects the data before harmonization, the HUH site moved much closer to the origin (i.e., grand average) and showed no marked separation from the other sites. This result indicates that the separation of the HUH site observed in Fig. 2 was caused by measurement bias, which was removed following harmonization. Furthermore, harmonization was effective in distinguishing patients and HCs scanned at the same site. Since patients with ASD were only scanned at the Showa University (SWA) site, the averages for patients with ASD (▲) and HCs (blue ●) scanned at this site were projected to nearly identical positions (Fig. 2). However, the two symbols are clearly separated from one another in Fig. 7. The effect of a psychiatric disorder (ASD) could not be observed in the first two PCs without harmonization but became detectable following the removal of measurement bias.

**Figure 7:**
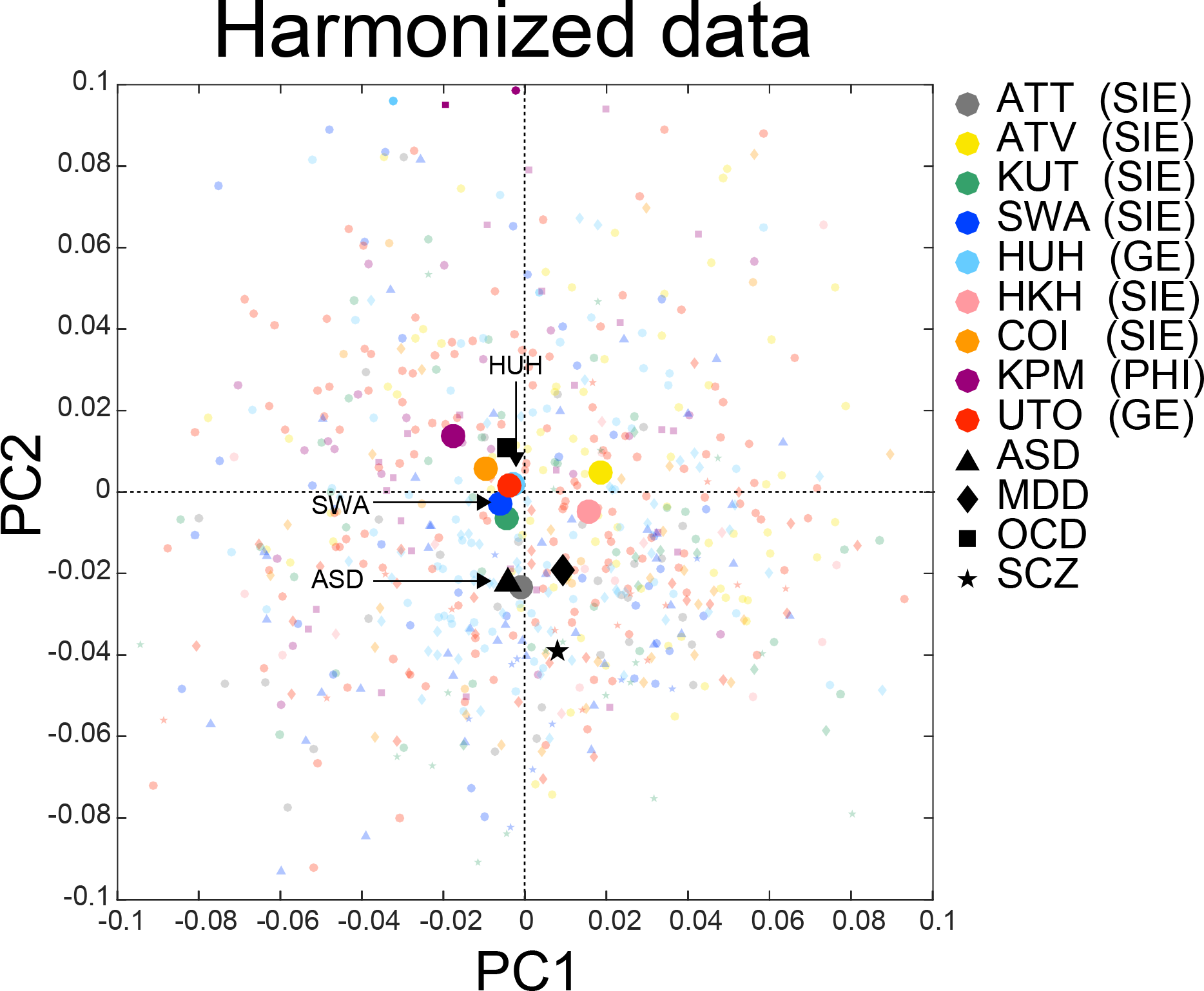
PCA dimension reduction in the SRPBS multi-disorder dataset after harmonization. All participants in the SRPBS multi-disorder dataset after harmonization projected into the first two principal components (PCs), as indicated by small, light-colored markers. The average across all healthy controls in each site and the average within each psychiatric disorder are depicted as dark-colored makers. The color of the marker represents the site, while the shape represents the psychiatric disorder. PCA: principal component analysis; SRPBS: Strategic Research Program for Brain Sciences; ATT: Siemens TimTrio scanner at Advanced Telecommunications Research Institute International; ATV: Siemens Verio scanner at Advanced Telecommunications Research Institute International; KUT: Siemens TimTrio scanner at Kyoto University; SWA: Showa University; HUH: Hiroshima University Hospital; HKH: Hiroshima Kajikawa Hospital; COI: Center of Innovation in Hiroshima University; KPM: Kyoto Prefectural University of Medicine; UTO: University of Tokyo; ASD: Autism Spectrum Disorder. MDD: Major Depressive Disorder. OCD: Obsessive Compulsive Disorder. SCZ: Schizophrenia. SIE: Siemens fMRI. GE: GE fMRI. PHI: Philips fMRI.

### Quantification of the effect of traveling-subject harmonization

To correct difference among sites there are three commonly used harmonization methods: (1) a ComBat method [16, 17, 19, 38], a batch-effect correction tool commonly used in genomics, site difference was modeled and removed; (2) a generalized linear model (GLM) method, site difference was estimated without adjusting for biological covariates (e.g., diagnosis) [16, 18, 22]; and (3) an adjusted GLM method, site difference was estimated while adjusting for biological covariates [16, 18] (see the “Harmonization procedures” in the Methods section). However, all these methods estimate the site difference without separating site difference into the measurement bias and the sampling bias and subtract the site difference from data. Therefore, existing harmonization methods might have pitfall to eliminate not only biologically meaningless measurement bias but also eliminate biologically meaningful sampling bias. Here, we tested whether the traveling-subject harmonization method indeed removes only the measurement bias and whether the existing harmonization methods simultaneously remove the measurement and sampling biases. Specifically, we performed 2-fold cross-validation evaluations in which the SRPBS multi-disorder dataset was partitioned into two equal-size subsamples (fold1 data and fold2 data) with the same proportions of sites. Between these two subsamples, the measurement bias is common, but the sampling bias is different (because the scanners are common, and participants are different). We estimated the measurement bias (or site difference including the measurement bias and the sampling bias for the existing methods) by applying the harmonization methods to the fold1 data and subtracted the measurement bias or site difference from the fold2 data. We then estimated the measurement bias in the fold2 data. For the existing harmonization methods, if the site difference estimated by using fold1 contains only the measurement bias, the measurement bias estimated in fold2 data after subtracting the site difference should be smaller than that of without subtracting the site difference (Raw). To separately estimate measurement bias and sampling bias in both subsamples while avoiding information leak, we also divided the traveling-subject dataset into two equal-size subsamples with the same proportions of sites and subjects. We concatenated one subsample of traveling-subject dataset to fold1 data to estimate the measurement bias for traveling-subject method (estimating dataset) and concatenated the other subsample of traveling-subject dataset to fold2 data for testing (testing dataset). That is, in the traveling-subject harmonization method, we estimated the measurement bias using the estimating dataset and removed the measurement bias from the testing dataset. By contrast, in the other harmonization methods, we estimated the site difference using the fold1 data (not including the subsample of traveling-subject dataset) and removed the site difference from the testing dataset. We then estimated the measurement bias using the testing dataset and evaluated the standard deviation of the magnitude distribution of measurement bias calculated in the same way as described in “Quantification of site differences” section. To verify whether important information such as participant factors and disorder factors are kept in the testing dataset, we also estimated the disorder factors and participant factors and calculated the ratio of the standard deviation of measurement bias to the standard deviation of participant factor and disorder factor as signal to noise ratios. This procedure was done again by exchanging the estimating dataset and the testing dataset (see the “2-fold cross-validation evaluation procedure” in the Methods section).

Fig. 8 shows that the standard deviation of measurement bias and the ratio of the standard deviation of measurement bias to the standard deviation of participant factor and disorder factor in the both fold data for the four harmonization methods and without harmonization (Raw). Our result shows that the reduction of the standard deviation of measurement bias from the Raw was highest in the traveling-subject method among all methods (29% reduction compared to 3% in the second highest value for ComBat method). Moreover, improvement in the signal to noise ratios were also highest in our method for participant factor (41% improvement compared to 3% in the second highest value for ComBat method) and for disorder factor (39% improvement compared to 3% in the second highest value for ComBat method). These results indicate that the traveling-subject harmonization method indeed removed the measurement bias and improved the signal to noise ratios.

**Figure 8:**
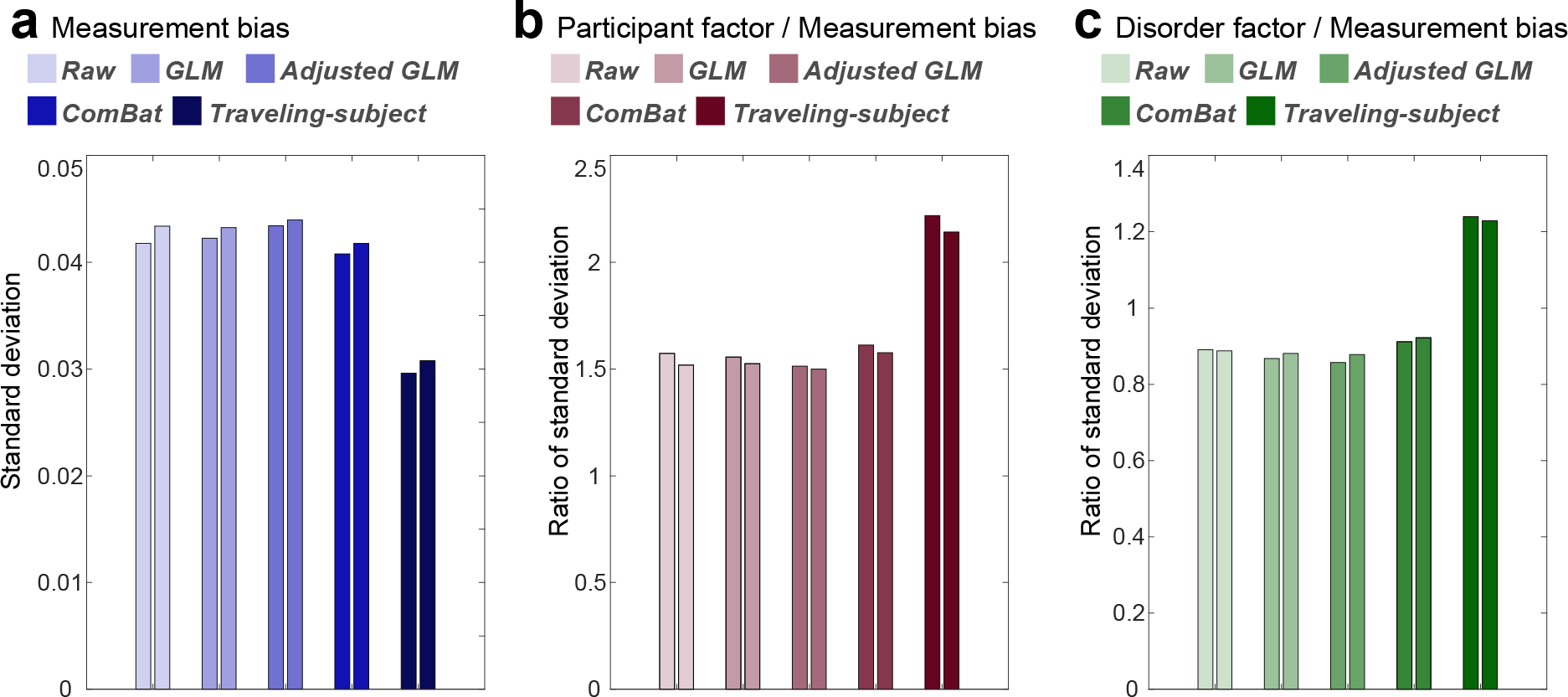
Reduction of the measurement bias and improvement of signal to noise ratios for different harmonization methods. (a) Standard deviation of the measurement bias. (b) Ratio of standard deviation of the measurement bias to standard deviation of the participant factor. (c) Ratio of standard deviation of the measurement bias to standard deviation of the disorder factor. Different colored columns show the results from different harmonization method. Two columns of the same color show the results of the two folds. GLM: generalized linear model.

## Discussion

In the present study, by acquiring a separate traveling-subject dataset and the SRPBS multi-disorder dataset, we separately estimated measurement and sampling biases for multiple sites, which we then compared with the magnitude of disorder factors. Furthermore, we investigated the origin of each bias in multi-site datasets. Finally, to overcome the problem of site difference, we developed a novel harmonization method that enabled us to subtract the measurement bias by using a traveling-subject dataset and achieved the reduction of the measurement bias by 29% and the improvement of the signal to noise ratios by 40%.

We assessed the effect sizes of measurement and sampling biases in comparison with the effects of psychiatric disorders on resting-state functional connectivity. Our findings indicated that measurement bias exerted significantly greater effects than disorder factors, whereas sampling bias was comparable to (or even larger than) the disorder effects (Fig. 3). However, we did not control for variations in disease stage and treatment in our dataset. Although controlling for such heterogeneity may increase the effect size of disorder factors, such control is not feasible when collecting big data from multiple sites. Therefore, it is important to appropriately remove measurement bias from heterogeneous patient data to identify relatively small disorder effects. This issue is essential for investigating the relationships among different psychiatric disorders because disease factors are often confounded by site differences. As previously mentioned, it is common for a single site to sample only a few types of psychiatric disorders (e.g., SCZ from site A and ASD from site B). In this situation, it is critical to dissociate disease factors from site differences. This dissociation can be accomplished by subtracting only the measurement bias which is estimated from traveling subject dataset.

Our results indicated that measurement bias is primarily influenced by differences in the phase-encoding direction, followed by differences in fMRI manufacturer, coil type, and scanner model (Fig. 5). These results are consistent with our finding of large measurement biases in the inferior brain regions (Fig. 4), the functional imaging of which is known to be influenced by the phase-encoding direction [28, 29]. Previous studies have reported that the effect because of the difference in the phase-encoding direction can be corrected using the field map obtained at the time of imaging [28, 39–41]. The field map was acquired in parts of the traveling-subject dataset; therefore, we investigated the effectiveness of field map correction by comparing the effect size of the measurement bias and the participant factor between functional images with and without field map correction. Our prediction was as follows: if field map correction is effective, the effect of measurement bias will decrease, while that of the participant factor will increase following field map correction. Field map correction using SPM12 (http://www.fil.ion.ucl.ac.uk/spm/software/spm12) reduced the effect of measurement bias in the inferior brain regions (whole brain: 3% reduction in the standard deviation of measurement bias) and increased the effect of the participant factor in the whole brain (3% increase in the standard deviation of the participant factor; Supplementary Figures 2a and 2b). However, the effect of measurement bias remained large in inferior brain regions (Supplementary Figure 2a), and hierarchical clustering analysis revealed that the clusters of the phase-encoding direction remained dominant (Supplementary Figure 2c). These results indicate that, even with field map correction, it is largely impossible to remove the influence of differences in phase-encoding direction on functional connectivity. Thus, harmonization methods are still necessary to remove the effect of these differences and other measurement-related factors. However, some distortion correction methods have been developed (e.g., top-up method and symmetric normalization) [42, 43], and further studies are required to verify the efficacy of these methods.

Our data supported the different-subpopulation model rather than the single-population model (Fig. 6), which indicates that sampling bias is caused by sampling from among different subpopulations. Furthermore, these findings suggest that, during big data collection, it is better to sample participants from several imaging sites than to sample many participants from a few imaging sites. These results were obtained only by combining the SRPBS multi-disorder database with a traveling-subject dataset (http://www.cns.atr.jp/decnefpro/). To the best of our knowledge, the present study is the first to demonstrate the presence of sampling bias in rs-fcMRI data, the mechanisms underlying this sampling bias, and the effect size of sampling bias on resting-state functional connectivity, which was comparable to that of psychiatric disorders. We analyzed sampling bias among HCs only, because the number of sites was too small to conduct an analysis of patients with psychiatric diseases.

We developed a novel harmonization method using a traveling-subject dataset (i.e., traveling-subject method), which was then compared with existing harmonization methods. Our results demonstrated that the traveling-subject method outperformed other conventional GLM-based harmonization methods and ComBat method. The traveling-subject method achieved reduction of the measurement bias by 29% compared to 3% in the second highest value for ComBat method and improvement of the signal to noise ratios by 40% compared to 3% in the second highest value for ComBat method. This result indicates that the traveling-subject dataset helps to properly estimate the measurement bias and also helps to harmonize the rs-fMRI data across imaging sites. To further quantitatively evaluate the harmonization method, we constructed biomarkers for psychiatric disorders based on rs-fcMRI data, which distinguishes between HCs and patients, and a regression model to predict participants’ age based on rs-fcMRI data using SRPBS multi-disorder dataset (see “Classifiers for MDD and SCZ, based on the four harmonization methods” and “Regression models of participant age based on the four harmonization methods” in Supplementary Information). We evaluated the generalization performance to independent validation dataset, which was not included in SRPBS multi-disorder dataset. The traveling-subject harmonization method improved the generalization performance of all these prediction models as compared with the case where harmonization was not performed. These results indicate that the traveling-subject dataset also helps the constructing a prediction model based on multi-site rs-fMRI data.

The present study possesses some limitations of note. The accuracy of measurement bias estimation may be improved by further expanding the traveling-subject dataset. This can be achieved by increasing the number of traveling participants or sessions per site. However, as mentioned in a previous traveling-subject study [20], it is costly and time-consuming to ensure that numerous participants travel to every site involved in big database projects. Thus, the cost-performance tradeoff must be evaluated in practical settings. The numbers of traveling participants and MRI sites used in this study (nine and 12, respectively) were larger than those used in a previous study (eight and eight, respectively) [20], and the number of total sessions in this study (411) was more than three times larger than that used in the previous study (128) [20]. Furthermore, although we estimated the measurement bias for each connectivity, hierarchical models of the brain (e.g., ComBat) may be more appropriate for improving the estimates of measurement bias.

In summary, by acquiring a separate traveling-subject dataset and the SRPBS multi-disorder database, we revealed that site differences were composed of biological sampling bias and engineering measurement bias. The effect sizes of these biases on functional connectivity were greater than or equal to the effect sizes of psychiatric disorders, highlighting the importance of controlling for site differences when investigating psychiatric disorders. Furthermore, using the traveling-subject dataset, we developed a novel traveling-subject method that harmonizes the measurement bias only by separating sampling bias from site differences. Our findings verified that the traveling-subject method outperformed conventional GLM-based harmonization methods and ComBat method. These results suggest that a traveling-subject dataset can help to harmonize the rs-fMRI data across imaging sites.

## Methods

### Participants

We used two resting-state functional MRI datasets for all analyses: (1) the SRPBS multi-disorder dataset, which encompasses multiple psychiatric disorders; (2) a traveling-subject dataset. The SRPBS multi-disorder dataset contains data for 805 participants (482 HCs from nine sites, 161 patients with MDD from five sites, 49 patients with ASD from one site, 65 patients with OCD from one site, and 48 patients with SCZ from three sites (Supplementary Table 1). Data were acquired using a Siemens TimTrio scanner at Advanced Telecommunications Research Institute International (ATT), a Siemens Verio scanner at Advanced Telecommunications Research Institute International (ATV), a Siemens Verio at the Center of Innovation in Hiroshima University (COI), a GE Signa HDxt scanner at HUH, a Siemens Spectra scanner at Hiroshima Kajikawa Hospital (HKH), a Philips Achieva scanner at Kyoto Prefectural University of Medicine (KPM), a Siemens Verio scanner at SWA, a Siemens TimTrio scanner at Kyoto University (KUT), and a GE MR750W scanner at the University of Tokyo (UTO). Each participant underwent a single rs-fMRI session for 5–10 min. The rs-fMRI data were acquired using a unified imaging protocol at all but three sites (Supplementary Table 2; http://www.cns.atr.jp/rs-fmri-protocol-2/). During the rs-fMRI scans, participants were instructed as follows, except at one site: “Please relax. Don’t sleep. Fixate on the central crosshair mark and do not think about specific things.” At the remaining site, participants were instructed to close their eyes rather than fixate on a central crosshair.

In the traveling-subject dataset, nine healthy participants (all male participants; age range, 24–32 years; mean age, 27±2.6 years) were scanned at each of 12 sites in the SRPBS consortium, producing a total of 411 scan sessions. Data were acquired at the sites included in the SRPBS multi-disorder database (i.e., ATT, ATV, COI, HUH, HKH, KPM, SWA, KUT, and UTO) and three additional sites: Kyoto University (KUS; Siemens Skyra) and Yaesu Clinic 1 and 2 (YC1 and YC2; Philips Achieva) (Supplementary Table 3). Each participant underwent three rs-fMRI sessions of 10 min each at nine sites, two sessions of 10 min each at two sites (HKH & HUH), and five cycles (morning, afternoon, next day, next week, next month) consisting of three 10-minute sessions each at a single site (ATT). In the latter situation, one participant underwent four rather than five sessions at the ATT site because of a poor physical condition. Thus, a total of 411 sessions were conducted [8 participants × (3×9＋2×2+5×3×1)+1 participant × (3×9＋2×2+4×3×1)]. During each rs-fMRI session, participants were instructed to maintain a focus on a fixation point at the center of a screen, remain still and awake, and to think about nothing in particular. For sites that could not use a screen in conjunction with fMRI (HKH & KUS), a seal indicating the fixation point was placed on the inside wall of the MRI gantry. Although we attempted to ensure imaging was performed using the same parameters at all sites, there were two phase-encoding directions (P→A and A→P), three MRI manufacturers (Siemens, GE, and Philips), four different numbers of coils (8, 12, 24, 32), and seven scanner types (TimTrio, Verio, Skyra, Spectra, MR750W, SignaHDxt, Achieva) (Supplementary Table 3).

All participants in all datasets provided written informed consent, and all recruitment procedures and experimental protocols were approved by the Institutional Review Boards of the principal investigators’ respective institutions (Advanced Telecommunications Research Institute International (ATR), Hiroshima University, Kyoto Prefectural University of Medicine, Showa University, The University of Tokyo).

### Preprocessing and calculation of the resting-state functional connectivity matrix

The rs-fMRI data were preprocessed using SPM8 implemented in MATLAB. The first 10 s of data were discarded to allow for T1 equilibration. Preprocessing steps included slice-timing correction, realignment, co-registration, segmentation of T1-weighted structural images, normalization to Montreal Neurological Institute (MNI) space, and spatial smoothing with an isotropic Gaussian kernel of 6 mm full-width at half-maximum. For the analysis of connectivity matrices, ROIs were delineated according to a 268-node gray matter atlas developed to cluster maximally similar voxels [26]. The BOLD signal time courses were extracted from these 268 ROIs. To remove several sources of spurious variance, we used linear regression with 36 regression parameters [44] such as six motion parameters, average signals over the whole brain, white matter, and cerebrospinal fluid. Derivatives and quadratic terms were also included for all parameters. A temporal band-pass filter was applied to the time series using a first-order Butterworth filter with a pass band between 0.01 Hz and 0.08 Hz to restrict the analysis to low-frequency fluctuations, which are characteristic of rs-fMRI BOLD activity [44]. Furthermore, to reduce spurious changes in functional connectivity because of head motion, we calculated frame-wise displacement (FD) and removed volumes with FD > 0.5 mm, as proposed in a previous study [45]. The FD represents head motion between two consecutive volumes as a scalar quantity (i.e., the summation of absolute displacements in translation and rotation). Using the aforementioned threshold, 5.4% ± 10.6% volumes (i.e., the average [approximately 13 volumes] ± 1 SD) were removed per 10 min of rs-fMRI scanning (240 volumes) in the traveling-subject dataset, 6.2% ± 13.2% volumes were removed per rs-fMRI session in the SRPBS multi-disorder dataset. If the number of volumes removed after scrubbing exceeded the average of –3 SD across participants in each dataset, the participants or sessions were excluded from the analysis. As a result, 14 sessions were removed from the traveling-subject dataset, 20 participants were removed from the SRPBS multi-disorder dataset. Furthermore, we excluded participants for whom we could not calculate functional connectivity at all 35,778 connections, primarily because of the lack of BOLD signals within an ROI. As a result, 99 participants were further removed from the SRPBS multi-disorder dataset.

### Principal component analysis

We developed bivariate scatter plots of the first two principal components based on a PCA of functional connectivity values in the SRPBS multi-disorder dataset (Fig. 2). To visualize whether most of the variation in the SRPBS multi-disorder dataset was still associated with imaging site after harmonization, we performed a PCA of functional connectivity values in the harmonized SRPBS multi-disorder dataset (Fig. 7). We used the traveling-subject method for harmonization, as described in the following section.

### Estimation of biases and factors

The participant factor (***p***), measurement bias (***m***), sampling biases (***s***_***hc***_, ***s***_***mdd***_, ***s***_***scz***_), and psychiatric disorder factor (***d***) were estimated by fitting the regression model to the functional connectivity values of all participants from the SRPBS multi-disorder dataset and the traveling-subject dataset. In this instance, vectors are denoted by lowercase bold letters (e.g., ***m***) and all vectors are assumed to be column vectors. Components of vectors are denoted by subscripts such as *m_k_*. To represent participant characteristics, we used a 1-of-K binary coding scheme in which the target vector (e.g., **x**_***m***_) for a measurement bias ***m*** belonging to site *k* is a binary vector with all elements equal to zero—except for element *k*, which equals 1. If a participant does not belong to any class, the target vector is a vector with all elements equal to zero. A superscript T denotes the transposition of a matrix or vector, such that **x**^T^ represents a row vector. For each connectivity, the regression model can be written as follows:

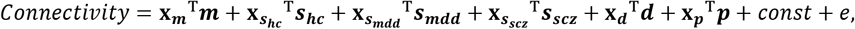

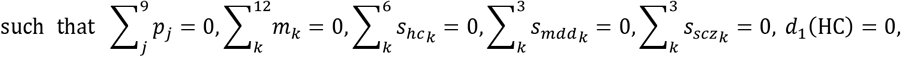

in which ***m*** represents the measurement bias (12 sites × 1), ***s***_***hc***_ represents the sampling bias of HCs (six sites × 1), ***s***_***mdd***_ represents the sampling bias of patients with MDD (three sites × 1), ***s***_***scz***_ represents the sampling bias of patients with SCZ (three sites × 1), ***d*** represents the disorder factor (3 × 1), ***p*** represents the participant factor (nine traveling subjects × 1), *const* represents the average functional connectivity value across all participants from all sites, and *e*~𝒩(0, γ^−1^) represents noise. For each functional connectivity value, we estimated the respective parameters using regular ordinary least squares regression with L2 regularization, as the design matrix of the regression model is rank-deficient. When regularization was not applied, we observed spurious anticorrelation between the measurement bias and the sampling bias for HCs, and spurious correlation between the sampling bias for HCs and the sampling bias for patients with psychiatric disorders (Supplementary Figure 3a, left). These spurious correlations were also observed in the permutation data in which there were no associations between the site label and data (Supplementary Figure 3a, right). This finding suggests that the spurious correlations were caused by the rank-deficient property of the design matrix. We tuned the hyper-parameter lambda to minimize the absolute mean of these spurious correlations (Supplementary Figure 3c, left).

### Analysis of contribution size

To quantitatively verify the magnitude relationship among factors, we calculated and compared the contribution size to determine the extent to which each bias type and factor explain the variance of the data in our linear mixed-effects model (Fig. 3d). After fitting the model, the *b*-th connectivity from subject *a* can be written, as follows:

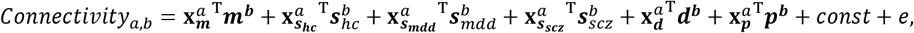

For example, the contribution size of measurement bias (i.e., the first term) in this model was calculated as

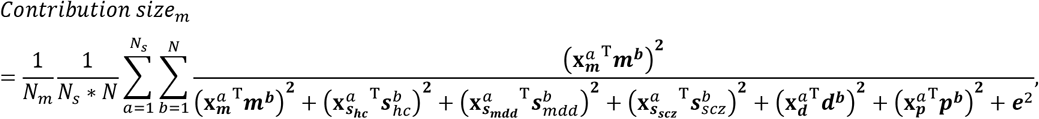

in which *N*_*m*_ represents the number of components for each factor, *N* represents the number of connectivities, *N*_*s*_ represents the number of subjects, and *Contribution size*_*m*_ represents the magnitude of the contribution size of measurement bias. These formulas were used to assess the contribution sizes of individual factors related to measurement bias (e.g., phase-encoding direction, scanner, coil, and fMRI manufacturer: Fig. 5b). We decomposed the measurement bias into these factors, after which the relevant parameters were estimated. Other parameters were fixed at the same values as previously estimated.

### Spatial characteristics of measurement bias, sampling bias, and each factor in the brain

To evaluate the spatial characteristics of each type of bias and each factor in the brain, we calculated the magnitude of the effect on each ROI. First, we calculated the median absolute value of the effect on each functional connection among sites or participants for each bias and participant factor. We then calculated the absolute value of each connection for each disorder factor. The uppercase bold letters (e.g., ***M***) and subscript vectors (e.g., ***m***_*k*_) represent the vectors for the number of functional connections:

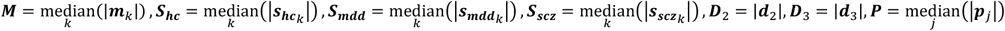

We next calculated the magnitude of the effect on ROIs as the average connectivity value between all ROIs, except for themselves.

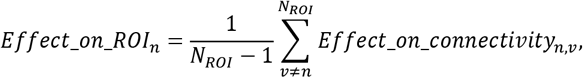

in which *N*_*ROI*_ represents the number of ROIs, *Effect_on_ROI*_*n*_ represents the magnitude of the effect on the *n*-th ROI, and *Effect_on_connectivity*_*n,v*_ represents the magnitude of the effect on connectivity between the *n*-th ROI and *v*-th ROI.

### Hierarchical clustering analysis for measurement bias

We calculated the Pearson’s correlation coefficients among measurement biases ***m***_*k*_ (*N* × 1, where *N* is the number of functional connections) for each site *k*, and performed a hierarchical clustering analysis based on the correlation coefficients across measurement biases. To visualize the dendrogram (Fig. 5), we used the “*dendrogram*”, “*linkage*”, and “*optimalleaforder*” functions in MATLAB (R2015a, Mathworks, USA).

### Comparison of models for sampling bias

We investigated whether sampling bias is caused by the differences in the number of participants among imaging sites, or by sampling from different populations among imaging sites. We constructed two models and investigated which model provides the best explanation of sampling bias. In the single-population model, we assumed that participants were sampled from a single population across imaging sites. In the different-population model, we assumed that participants were sampled from different populations among imaging sites. We first theorized how the number of participants at each site affects the variance of sampling biases across connectivity values, as follows:

In the *single-population model*, we assumed that the functional connectivity values of each participant were generated from an independent Gaussian distribution, with a mean of 0 and a variance of *ξ*^2^ for each connectivity value. Then, the functional connectivity vector for participant *j* at site *k* can be described as

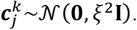

Let ***c***_*k*_ be the vector of functional connectivity at site *k* averaged across participants. In this model, ***c***_*k*_ represents the sampling bias and can be described as

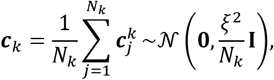

in which *N*_*k*_ represents the number of participants at site *k*. The variance across functional connectivity values for ***c***_*k*_ is described as

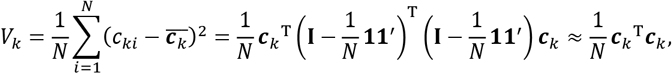

in which **1** represents the *N* × 1 vector of ones and **I** represents the *N* × *N* identity matrix. Since *N* equals 35,778 and 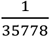 is sufficiently smaller than 1, we can approximate

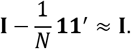

Then, the expected value of variance across functional connectivity values for sampling-bias can be described as

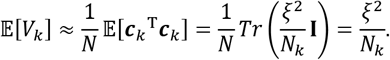

In the different-population model, we assumed that the functional connectivity values of each participant were generated from a different independent Gaussian distribution, with an average of ***β***_***k***_ and a variance of *ξ*^2^ depending on the population of each site. In this situation, the functional connectivity vector for participant *j* at site *k* can be described as

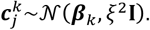

Here, we assume that the average of the population ***β***_***k***_ is sampled from an independent Gaussian distribution with an average of 0 and a variance of σ^2^. That is, ***β***_*k*_ is expressed as

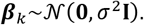

The vector of functional connectivity for site *k* averaged across participants can then be described as

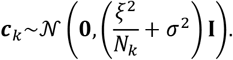

The variance across functional connectivity values for ***c***_*k*_ can be described as

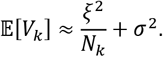

In summary, the variance of sampling bias across functional connectivity values in each model is expressed by the number of participants at a given site, as follows:

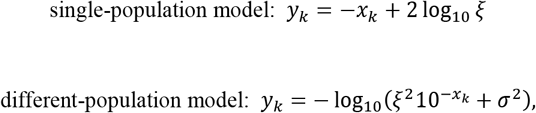

in which *y*_*k*_ = log_10_(*v*_*k*_), *v*_*k*_ represents the variance across functional connectivity values for 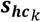, 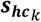, represents the sampling bias of HCs at site *k* (*N* × 1: *N* is the number of functional connectivity), *x*_*k*_ = log_10_(*N*_*k*_), and *N*_*k*_ represents the number of participants at site *k*. We estimated the parameters *ξ* and σ using the MATLAB (R2015a, Mathworks, USA) optimization function “*fminunc*”. To simplify statistical analyses, sampling bias was estimated based on functional connectivity in which the average across all participants was set to zero.

We aimed to determine which model provided the best explanation of sampling bias in our data by calculating the corrected Akaike information criterion (AICc; under the assumption of a Gaussian distribution) for small-sample data [36, 37], as well as BIC:

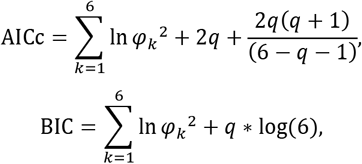

in which 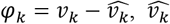 represents the estimated variance, and *q* represents the number of parameters in each model (1 or 2).

To investigate prediction performance, we used leave-one-site-out-cross-validation in which we estimated the parameters *ξ* and σ using data from five sites. The variance of sampling bias was predicted based on the number of participants at the remaining site. This procedure was repeated to predict variance values for sampling bias at all six sites. We then calculated the absolute errors between predicted and actual variances for all sites.

### Harmonization procedures

We compared four different harmonization methods for the removal of site differences, as described in the main text.

#### Traveling-subject harmonization

Measurement biases were estimated by fitting the regression model to the combined SRPBS multi-disorder and traveling-subject datasets in the same way in “Estimation of biases and factors” section. For each connectivity, the regression model can be written as follows:

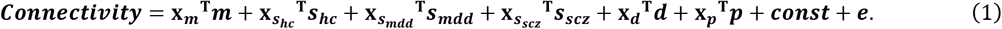

Measurement bias were removed by subtracting the estimated measurement biases. Thus, the harmonized functional connectivity values were set, as follows:

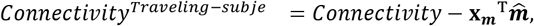

in which 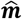 represents the estimated measurement bias.

#### GLM harmonization

The GLM harmonization method adjusts the functional connectivity value for site difference using GLM. Site differences were estimated by fitting the regression model, which included site label only, to the SRPBS multi-disorder dataset only. The regression model can be written as

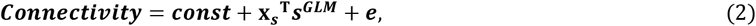

in which ***s***^*GLM*^ represents the site difference (nine sites × 1). For each functional connectivity value, we estimated the parameters using regular ordinary least squares regression. Site differences were removed by subtracting the estimated site differences. Thus, the harmonized functional connectivity values were set, as follows:

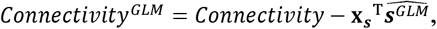

in which 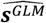 represents the estimated site difference.

#### Adjusted GLM harmonization

Site differences were estimated by fitting the regression model, which included site label and diagnosis label, to the SRPBS multi-disorder dataset. The regression model can be written as

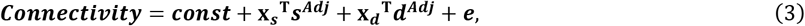

In which ***s***^*Adj*^ represents the site difference (nine sites × 1). For each functional connectivity value, we estimated the parameters via regular ordinary least squares regression. Site differences were removed by subtracting the estimated site difference only. Thus, the harmonized functional connectivity values were set, as follows:

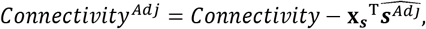

in which 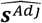 represents the estimated site difference.

#### ComBat harmonization

The ComBat harmonization model [16, 17, 19, 38] extends the adjusted GLM harmonization method in two ways: (1) it models site-specific scaling factors and (2) it uses empirical Bayesian criteria to improve the estimation of site parameters for small sample sizes. The model assumes that the expected connectivity value can be modeled as a linear combination of the biological variables and the site differences in which the error term is modulated by additional site-specific scaling factors.

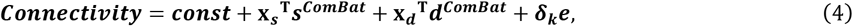

in which ***s***^*ComBat*^ represents the site difference (nine sites × 1), and *δ*_*k*_ represents the scale parameter for site differences at site *k* for the respective connectivity value. The harmonized functional connectivity values were set, as follows:

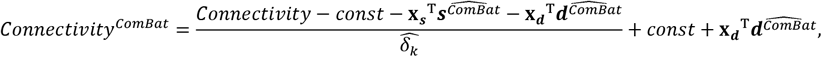

in which 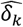, 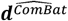, and 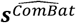 are the empirical Bayes estimates of *δ*_*k*_, ***d***^*ComBat*^, and ***s***^*ComBat*^, respectively using “combat” function in http://github.com/Jfortin1/ComBatHarmonization. Thus, ComBat simultaneously models and estimates biological and nonbiological terms and algebraically removes the estimated additive and multiplicative site differences. Of note, in the ComBat model, we included diagnosis as covariates to preserve important biological trends in the data and avoid overcorrection.

### 2-fold cross-validation evaluation procedure

We compared four different harmonization methods for the removal of site difference or measurement bias by 2-fold cross-validation, as described in the main text. In the traveling-subject harmonization method, we estimated the measurement bias by applying the regression model written in equation (1) in “Harmonization procedures” section to the estimating dataset. Thus, the harmonized functional connectivity values in testing dataset were set, as follows:

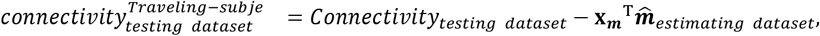

in which 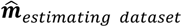 represents the estimated measurement bias using the estimating dataset.

By contrast, in the other harmonization methods, we estimated the site differences by applying the regression models written in equations (2)–(4) in “Harmonization procedures” section to the estimating dataset (fold1 data). Thus, the harmonized functional connectivity values in testing dataset were set, as follows:

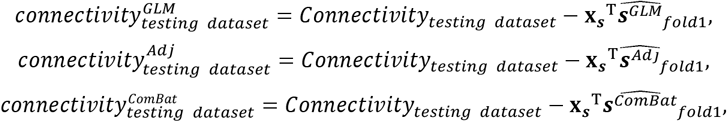

in which 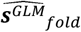, 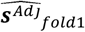, 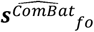 represents the estimated site differences using fold1 data.

We then estimated the measurement bias, participant factor, and disorder factors by applying the regression model written in equation (1) to the harmonized functional connectivity values in the testing dataset. Finally, we evaluated the standard deviation of the magnitude distribution of measurement bias calculated in the same way as described in “Quantification of site differences” section among the harmonization methods. This procedure was done again by exchanging the estimating dataset and the testing dataset.

## Code availability

All codes used for the analyses are available from the authors on request.

## Data availability

All relevant data are available from the authors on request. All data can be downloaded publicly from the following site: http://bicr-resource.atr.jp/decnefpro/.

## Acknowledgements

This study was conducted under the “Development of BMI Technologies for Clinical Application” of the Strategic Research Program for Brain Sciences, and the contract research Grant Number JP18dm0307008 supported by the Japan Agency for Medical Research and Development (AMED). This study was also partially supported by the ImPACT Program of the Council for Science, Technology and Innovation (Cabinet Office, Government of Japan). H.I. was partially supported by JSPS KAKENHI 26120002.

## Author contributions

A.Y., N.Y., and H.I. designed the study. N.Y., T.I., T.Y., N.I., M.T., Y.Y., A.K., N.O., T.Y., K.M., R.H., G.O., Y.S., J.N., Y.S., K.K., N.K., H.T., Y.O. and S.T. recruited participants of the study, collected their clinical and imaging data and constructed the database. A.Y. performed data preprocessing and data analysis under the super vision of G.L., J.M., O.Y., M.K., and H.I., and A.Y., O.Y., M.K. and H.I. primarily wrote the manuscript.

## Competing financial interests

The authors declare no competing financial interests.

